# Catecholaminergic cell type-specific expression of Cre recombinase in knock-in transgenic rats generated by the *Combi*-CRISPR technology

**DOI:** 10.1101/2022.04.13.488137

**Authors:** Natsuki Matsushita, Kayo Nishizawa, Shigeki Kato, Yoshio Iguchi, Ryoji Fukabori, Kosei Takeuchi, Yoshiki Miyasaka, Tomoji Mashimo, Kazuto Kobayashi

**Author notes:** Both authors contributed equally to this work (Natsuki Matsushita, Kayo Nishizawa). **Corresponding author:** Kazuto Kobayashi, Ph.D., Department of Molecular Genetics, Institute of Biomedical Sciences, Fukushima Medical University School of Medicine, Fukushima 960-1295, Japan.

## Abstract

**Background:** Cell groups containing catecholamines provide a useful model to study the molecular and cellular mechanisms underlying the morphogenesis, physiology, and pathology of the central nervous system. For this purpose, it is necessary to establish a system to induce catecholaminergic group-specific expression of Cre recombinase. Recently, we introduced a gene cassette encoding 2A peptide fused to Cre recombinase into the site between the C-terminus and translational termination codons of the rat tyrosine hydroxylase (TH) open reading frame by the *Combi*-CRISPR technology, which is a genomic editing method to enable an efficient knock-in (KI) of long DNA sequence into a target site. However, the expression patterns of the transgene and its function as well as the effect of the mutation on the biochemical and behavioral phenotypes in the KI strains have not been characterized yet.

**New Method:** We aimed to evaluate the usefulness of TH-Cre KI rats as an experimental model for investigating the structure and function of catecholaminergic neurons in the brain.

**Results:** We detected cell type-specific expression of Cre recombinase and site-specific recombination activity in the representative catecholaminergic groups in the TH-Cre KI rat strains. In addition, we measured TH expression level and catecholamine accumulation in the brain regions, and spontaneous locomotion, indicating that catecholamine metabolism and general behavior are apparently normal in these KI rats.

**Conclusions:** TH-Cre KI rat strains produced by the *Combi*-CRISPR system offer a beneficial model to study the molecular and cellular mechanics for the morphogenesis, physiology, and pathology of catecholamine-containing neurons in the brain.

## 1. Introduction

Cell groups containing catecholamines including dopamine (DA), norepinephrine (NE), and epinephrine (EPI) are localized in diverse regions of the central nervous system (Björklund and Dunnett, 2007; Robertson et al., 2013) and play important roles in a wide range of brain functions, such as motor control, cognition, attention, learning, emotion, and endocrine regulation (Ranjbar-Slamloo and Fazlali, 2019; Robbins and Everitt, 1982). Therefore, dysfunctions in catecholamine-containing neurons and catecholamine transmission are implicated in the pathogenesis of various neurological and neuropsychiatric disorders. For instance, degeneration of nigrostriatal DA neurons causes Parkinson’s disease characterized mainly by bradykinesia, rigidity, and tremor (Berardelli et al., 2013; Schapira et al., 2009). Abnormalities in the mesolimbic or mesocortical DA transmission are related to symptoms of schizophrenia, attention-deficit hyperactivity disorder, and substance use disorder (Cassidy et al., 2018; Nestler and Lüscher, 2019; Silvetti et al., 2013). In addition, deficiency of NE transmission results in reduced arousal level, depression, fatigue, and sleep disturbance (Mäki-Marttunen et al., 2020; Ross and Van Bockstaele, 2020), whereas excessive NE function is associated with anxiety, agitation, and restlessness (McCall et al., 2015).

Catecholamine-containing neurons provide a model system to investigate molecular mechanisms underlying the development and survival of specific neuronal types in the brain. DA neurons in the ventral midbrain are generated through signal pathways originating from the floor plate and midbrain-hindbrain boundary and a network of multiple transcription factors, such as Lmx1a/b, Nurr1, and Pitx3 (Arenas et al., 2015; Wang et al., 2020). Generation of NE neurons in the hindbrain depends on the roles of some morphologic proteins and transcription factors (Hirsch et al., 1998; Morin et al., 1997; Pattyn et al., 2000; Vogel-Höpker and Rohrer, 2002). In addition, survival of midbrain DA neurons is promoted through multiple neurotrophic factors, including brain-derived neurotrophic factor, glial cell line-derived neurotrophic factor, and cerebral DA neurotrophic factor (Jaumotte et al., 2021; Kumar et al., 2015; Palasz et al., 2020), although studies on survival factors for central NE neurons have been limited (Holm et al., 2002; Reiriz et al., 2002). In particular, a better understanding of the mechanisms for development and survival of midbrain DA neurons provides a useful approach for clinical treatments with cell replacement and gene therapy for Parkinson’s disease (Buttery and Barker, 2020).

To study the molecular and cellular mechanics for morphogenesis, physiology, and pathology of catecholamine-containing neurons, it is necessary to establish a system for cell type-specific expression of genes of interest. One promising approach is the use of Cre-*lox*P-mediated recombination system (Gerfen et al., 2013; Witten et al., 2011). Transgenic mouse and rat lines were generated that express Cre recombinase by using a short DNA fragment or bacterial artificial chromosome (BAC) clone containing a promoter region of tyrosine hydroxylase (TH) gene (Gelman et al., 2003; Gong et al., 2007; Witten et al., 2011). However, these conventional methods are known to produce occasional discrepancies in the cell type specificity of transgene expression from endogenous expression patterns of the target gene, depending on the integrated position and copy number of the transgene (Gong et al., 2007; Matsushita et al., 2004). As compared to these methods, a knock-in (KI) strategy, in which the transgene is inserted into the targeted gene locus by using gene targeting with embryonic stem cells or genomic editing, is more useful to achieve the cell type specificity of transgene expression. Indeed, KI mouse and rat lines were generated with targeted insertion of Cre transgene into the 3’-untralslated region of TH gene mediated by an internal ribosomal entry site (IRES) (Brown et al., 2013; Lindeberg et al., 2004). However, some previous studies have reported that the bicistronic structure using the IRES shows difficulties in the efficient expression of the reporter gene downstream of the IRES (Hennecke et al., 2001; Zhou et al., 1998). In addition, the insertion of the IRES is reported to affect the expression level of the target gene products, and the TH-IRES-Cre KI mutation results in a mild reduction of TH expression level (Brown et al., 2013).

In contrast, the CRISPR-Cas9 system has been used for KI mutagenesis by using homology-directed repair with a donor DNA template (Inui et al., 2014; Yang et al., 2013). However, using this technology it is difficult to introduce efficiently long DNA sequences into the target site (Miura et al., 2015; Quadros et al., 2017). Recently, an efficient KI strategy termed *Combi*-CRISPR has been established on the basis of both non-homologous end-joining and homology-directed repair pathways (Yoshimi et al., 2021). This strategy was used to introduce a gene cassette containing 2A peptide connected to Cre transgene into the site between the C-terminus and translational termination codons of rat TH open reading frame, and two kinds of KI founders were obtained carrying an artificial insertion of a single nucleotide, A or T, into intron 12 of the TH gene (Yoshimi et al., 2021). However, the expression patterns of the transgene and its function in these two KI strains, as well as the effect of the mutations on the biochemical and behavioral phenotypes in the animals, have not been characterized yet.

In the present study, we examined the expression patterns of Cre recombinase in the representative DA and NE cell groups in the two TH-Cre KI rat strains, showing similar patterns of the transgene expression between these strains. Using one of the two strains, we then analyzed the transgene expression patterns in other catecholamine-containing cell groups, as well as recombinase activity in the representative neurons. In addition, we measured TH expression level and catecholamine accumulation in the brain regions of the KI rats, and monitored their spontaneous behavior in the open field, indicating that catecholamine metabolism and general activity in these rats are not apparently influenced by the KI mutation. Therefore, TH-Cre KI rat strains produced by the *Combi*-CRISPR system provide a powerful tool to study the molecular and cellular mechanisms underlying the morphogenesis, physiology, and pathology of brain catecholamine-containing neurons.

## 2. Materials and methods

### 2.1. Animals

Animal care and handling procedures were conducted in accordance with the guidelines established by the Laboratory Animal Research Center of Fukushima Medical University. All procedures were approved by the Fukushima Medical University Institutional Animal Care and Use Committee. Rats were maintained at 22 ± 2°C and 60% humidity in a 12-hr light/12-hr dark cycle, and food and water were continuously available. We made all efforts to minimize the number of animals used, and their suffering. Forty rats (8–12 weeks old) were used for the present study.

TH-Cre KI transgenic rats (KI/2-1 and KI/2-2 strains) were generated by using *Combi*-CRISPR technology as described previously (Yoshimi et al., 2021). In these strains, a gene cassette encoding 2A peptide (Kim et al., 2011) fused to Cre recombinase (GenBank accession number YP006472; Łobocka et al., 2004) with the nuclear localization signal followed by the polyadenylation signal from the bovine growth hormone gene (GenBank accession number M57764; Gordon et al., 1983) was introduced between the C-terminus of protein-coding region and 3’-untranslated region in exon 13 of the rat TH gene (GenBank accession number M23598; Brown et al., 1987), and a single nucleotide T (for KI/2-1 strain) or A (for KI/2-2 strain) was introduced in intron 12 of the rat TH gene during the process of genomic editing (see Supplementary Fig. 1 A, B). TH-Cre KI transgenic rats were mated with wild-type (WT) Long Evans rats (Charles River Laboratories, Wilmington, MA) to obtain offspring. Genotyping of the offspring was carried out with polymerase chain reaction (PCR) with genomic DNA prepared from tail clips (see Supplementary Fig. 1C).

Primers used were 5’-GGGCTTTAGTCTCCTGAATGTC-3’ for the forward primer and 5’-CTTTGTGGTGACGCAGTGACCAG-3’ for the reverse primer. The amplifications were performed with 25 cycles of denaturation at 96 °C for 10 s, annealing at 58 °C for 20 s, and extension at 72 °C for 75 s. The KI/2-1 and KI/2-2 strains were identified by nucleotide sequencing of the corresponding region in intron 12 by using by the capillary electrophoresis instrument (3500 Genetic Analyzer with a Data Collection Software; Thermo Fisher Scientific, Waltham, MA).

### 2.2. Viral vector preparation

Adeno-associated viral (AAV) vector serotype 2 was prepared using an AAV Helper-Free system (Agilent Technologies, Santa Clara, CA) as described previously (Kato et al., 2018). HEK293T cells (American Type Culture Collection, Manassas, VA) were transfected with the plasmids encoding transfer gene, adeno-helper gene, and adenoviral genes required for AAV replication and encapsulation through the calcium phosphate precipitation method. In the present study, as the flip/excision switch (FLEX) system we used the transfer plasmid containing the gene cassette encoding green fluorescent protein (GFP; GenBank accession number L29345; Inouye and Tsuji, 1994) in the inverted orientation, which is flanked by a pair of double recognition sites consisting of *lox*P and *lox*2272 sites in the opposite direction with an alternate order at both ends of the cassette. Another gene cassette encoding TurboFP635 (GenBank accession number AB982101; Takai et al., 2015) was inserted between two recognition sites at the 3’-end of the GFP sequence to protect non-specific expression of transgene in the tissues after the injection of AAV-FLEX-GFP vector (Matsushita et al., 2022 [bioRxiv]). The crude viral lysate was purified with two rounds of CsCl gradient ultracentrifugation at 100,000×*g* and 4°C for 23 h using 13.5-mL Quick-Seal tubes and a NVT65 rotor (Beckman Coulter, Brea, CA). Each tube was placed gently and 500– 750 µL of white turbid bands were collected. The solution was applied to three rounds of dialysis against 1 liter of PBS by using a Slide-A-Lyzer dialysis cassette (MWCO10,000; Thermo Fisher Scientific). The dialyzed solution was finally concentrated by centrifugation through a Vivaspin Turbo filter (Sartorius, Göttingen, Germany). The viral genome titer was determined by quantitative PCR using the TaqMan system (Thermo Fisher Scientific). PCR was carried out on duplicate samples by using a StepOne real-time PCR system (Applied Biosystems, Tokyo, Japan) under the following conditions: one cycle of 95°C for 3 min, followed by 40 cycles of 95°C for 15 s and 60°C for 30 s. The standard curve was prepared on the basis of serial dilutions of viral DNA control templates ranging from 4 × 10^4^ to 4 × 10^7^ genome copies/mL.

### 2.3. Viral vector injection

TH-Cre KI/2-1 rats were anaesthetized with isoflurane (4% induction and 1.5% maintenance), and AAV-FLEX-GFP vector (3.7 × 10^12^ genome copies/mL) was introduced into the brain regions, including the ventral tegmental area (VTA; 0.5 µL/site × 6 sites), substantia nigra pars compacta (SNc; 0.5 µL/site × 4 sites), and locus coeruleus (LC; 0.5 µL/site × 4 sites) through a glass microinjection capillary connected to a microinfusion pump (ESP-32; Eicom, Kyoto, Japan) at a constant rate of 0.1 μL/min according to the following coordinates of the rat brain (Paxinos and Watson, 2005). The respective anteroposterior (AP), mediolateral (ML), and dorsoventral (DV) coordinates from the bregma and dura (mm) for VTA injection were −5.0/0.8/−8.1 (Site 1), −5.0/−0.8/−8.1 (Site 2), −5.4/0.6/−8.0 (Site 3), −5.4/−0.6/−8.0 (Site 4), −5.9/0.4/−7.9 (Site 5), and −5.9/−0.4/−7.9 (Site 6). The respective AP/ML/DV coordinates for SNc injection were −5.0/2.5/−7.7 (Site 1), −5.0/−2.5/−7.7 (Site 2), −5.4/2.3/−7.3 (Site 3), and −5.4/−2.3/ −7.3 (Site 4). The respective coordinates for LC injection were −9.70/1.2/−5.3 (Site 1), −9.70/−1.2/−5.3 (Site 2), −9.90/0.2/−5.3 (Site 3), and −9.90/−0.2/−5.3 (Site 4).

### 2.4. Immunohistochemical staining

Rats were anesthetized with sodium pentobarbital (100 mg/kg, i.p.) and perfused transcardially with PBS, followed by 4% paraformaldehyde in 0.1 M phosphate buffer (pH 7.4). For immunofluorescence histochemistry, fixed brains were cut into sections (30-μm thick) through a coronal plane with a cryostat. Sections were incubated with rabbit anti-TH antibody (1:1000; Millipore, Billerica, MA) and mouse anti-Cre antibody (1:1000; Millipore, Billerica, MA) or goat anti-GFP antibody (1 μg/mL; Frontier Science, Hokkaido, Japan). Alexa Fluor 488-conjugated donkey anti-rabbit antibody (1:1000; Thermo Fisher Scientific, Waltham, MA), Cy3-conjugated donkey anti-mouse antibody (1:1000; Jackson ImmunoResearch Laboratories, West Grove, PA), and Cy3-conjugated donkey anti-goat antibody (1:1000; Jackson ImmunoResearch Laboratories) were used as species-specific secondary antibodies for detecting TH, Cre, and GFP immunoreactive signals, respectively. For measurement of fluorescent intensity to quantify TH protein expression, fixed brains were cut into sections (50-μm thick) with a vibrating blade slicer (DTK-1000, Dosaka EM Co., Kyoto, Japan). Sections were incubated with mouse anti-TH antibody (1:450; Millipore) and goat anti-microtubule associated protein 2 (MAP2) antibody (1:200; Frontier Science). Alexa Fluor 488-conjugated donkey anti-mouse antibody (1:200; Jackson ImmunoResearch Laboratories) and Cy3-conjugated donkey anti-goat antibody (1:200; Jackson ImmunoResearch Laboratories) were used as species-specific secondary antibodies for detecting TH and MAP2 immunoreactive signals, respectively. Fluorescent images were obtained with a confocal laser-scanning microscope (Nikon A1, Tokyo, Japan) equipped with proper filter cube specifications.

For cell counts, we selected two sections for each brain region, VTA, SNc, LC, olfactory bulb (OB), medial zona incerta (MZI), mediobasal hypothalamus (MBH), and nucleus tract solitaries (NTS), from each of the six animals to set regions of interest (ROIs; 0.6 × 0.6 mm). For double immunostaining against TH and Cre in the KI rats, the number of immuno-positive cells in the ROIs was counted. The ratio of the number of Cre^+^ and TH^+^ neurons relative to the total number of TH^+^ neurons were calculated. For double staining against TH and GFP in the vector-injected rats, ROIs were set in the VTA, SNc, and LC, and the number of stained cells in the ROIs was counted. The ratio of the number of GFP^+^ and TH^+^ neurons relative to the total number of TH^+^ neurons were calculated.

For measurement of fluorescent intensity, confocal images were quantified using Fiji/Image J (Schindelin et al., 2012). Each stack was split to the respective green (TH) and red (MAP2) image components and blurred by a Gaussian function to attenuate noise. For detection and segmentation of TH^+^ somas, the image was binarized by intensity thresholding using the Max Entropy algorithm or Trainable Weka Segmentation. The soma area information was redirected to the MAP2 component, and the mean gray value for each soma area was measured in each component. We detected an average of 62.6 (SNc), 59.8 (VTA), and 68.1 (LC) somas per animal. The ratio of TH/MAP2 intensity was calculated for each TH^+^ soma, and averages per animal were calculated for statistical analysis.

### 2.5. Catecholamine analysis

DA and NE contents were determined as described previously (Kobayashi et al., 1994) with some modifications. Rats were anesthetized with 3% isoflurane and decapitated for tissue preparation. The anterior cingulate cortex (ACC), nucleus accumbens (NAc) and dorsal striatum (DS) were dissected and homogenized immediately in 0.2 M perchloric acid containing 0.1 mM EDTA and the proper concentration of isoproterenol. After the tissue homogenates had been centrifuged, the pH of the supernatant was adjusted to 3.0 by adding 1 M sodium acetate. The samples were filtered and injected into the high-performance liquid chromatography system (SC-5ODS, 3.0 mm × 150 mm; Eicom, Kyoto, Japan) with the mobile phase containing 15% methanol in 0.1 M sodium citrate, 0.1 M citric acid, 0.5 mM sodium octanesulfonate, and 0.15 mM EDTA (pH 3.5) equipped with an electrochemical detector (ECD-300, Eicom). The detector potential was maintained at 750 mV versus the Ag/AgCl electrode.

### 2.6. Behavioral analysis

The open field consists of a gray plastic board (90 × 90 cm) surrounded by gray walls (45 cm in height). Rats were placed in the arena and allowed to explore it for 60 min. Their behaviors were recorded and analyzed using a video tracking software (Viewer2; Biobserve GmbH, Bonn, Germany). The distance traveled in the entire arena was analyzed in blocks of 10 min. In addition, two zones were defined, the central zone of the arena (0.36 m^2^) and a peripheral zone adjacent to the walls (a 15-cm wide corridor-like part, 0.45 m^2^), and distance traveled in each zone was determined.

### 2.7. Statistical analysis

Although no statistical methods were used to predetermine the sample size for each measure, we employed similar sample sizes to those that were reported in previous publications from our labs, which are generally accepted in the field. Unpaired *t*-tests and two-way analyses of variance (ANOVAs) were performed in a two-tailed manner using SPSS ver. 25 (IMB Corp., Armonk, NY). The reliability of the results was assessed against a type I error (*α*) of 0.05. Prior to the unpaired *t*-test, we assessed equality of variances for two groups with Levene’s test, and if this was not the case, we reported the statistics (*F* and *p*) and then corrected the degree of freedom according to Welch’s method.

## 3. Results

### 3.1. Cre expression in catecholaminergic cell types in KI transgenic rat strains

We examined the expression patterns of Cre recombinase in catecholamine-containing neurons in TH-Cre KI/2-1 and KI/2-2 rat strains, and compared them between the two strains. We selected A10 and A9 cell groups in the VTA and SNc, respectively, as representatives of DA neurons as well as A6 cell group in the LC as a representative of NE neurons. Coronal sections from the VTA, SNc, and LC were prepared from the brains of TH-Cre KI/2-1 and KI/2-2 strains (Fig. 1A). The sections were stained by double immunohistochemistry with anti-TH and anti-Cre antibodies (see Fig. 1B for typical images). Cre^+^ signals were found in the vast majority of the TH^+^ neurons in the VTA (97% ± 0.8% for KI/2-1 and 94% ± 2.0% for KI/2-2) and the SNc (95% ± 0.8% for KI/2-1 and 94% ± 0.8% for KI/2-2) (n = 6 animals), showing no significant difference in the frequency of Cre transgene expression in each brain region between the two rat strains (unpaired *t*-tests, *t*_10_ = 1.501, *p* = 0.164 for the VTA, and *t*_10_ = 0.622, *p* = 0.548 for the SNc). The signals were also observed in a number of TH^+^ neurons in the LC (100% for both KI/2-1 and KI/2-2) (n = 6 animals), indicating the equivalent frequency of Cre expression between the two strains. These results suggest that a single nucleotide insertion (T or A) in intron 12 of the rat TH gene does not change the expression patterns of Cre recombinase in the brains of TH-Cre KI transgenic rats. Therefore, we used the TH-Cre KI/2-1 strain for the following histological, biochemical, and behavioral analyses.

**Figure 1.**
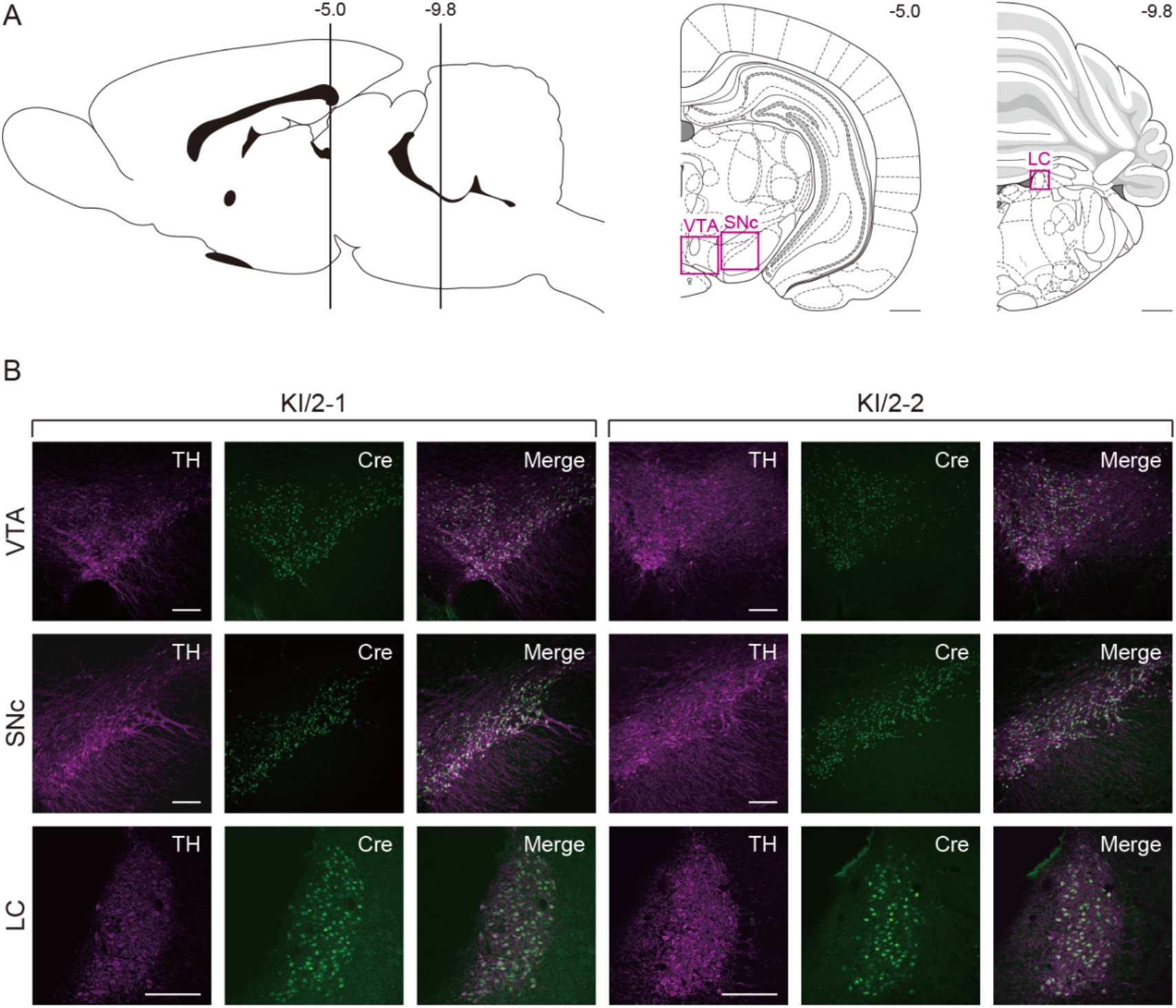
Expression of Cre transgene in the ventral midbrain and hindbrain of two KI transgenic rat strains. (A) Representative sections from the VTA, SNc, and LC used for double fluorescent immunohistochemistry with anti-TH and anti-Cre antibodies. Anteroposterior coordinates from the bregma (mm) are shown. (B) Double immunostaining for TH and Cre. Sections from the SNc, VTA, and LC were prepared from the TH-Cre KI/2-1 and KI/2-2 strains, and doubly immunostained with two kinds of antibodies. TH-positive, Cre-positive, and merged images are shown. Scale bar: 1 mm (A), 200 µm (B).

We next investigated the expression patterns of Cre recombinase in other catecholamine-containing neurons in the brain of TH-Cre KI/2-1 transgenic rats. We chose A16 cell group in the OB, A13 cell group in the MZI, and A12 cell group in the MBH as representatives of DA neurons as well as A2/C2 cell group in the NTS as representatives of NE/EPI neurons. Coronal sections from the OB, MZI, MBH, and NTS were prepared from the brains of the TH-Cre KI/2-1 strain (Fig. 2A). The sections were stained by double immunohistochemistry for TH and Cre (see Fig. 2B for typical images). Cre^+^ signals were observed in a number of TH^+^ neurons in the OB (99% ± 0.1%), MZI (99% ± 0.2%), MBH (100%), and NTS (100%) (n = 6 animals). These results indicate cell type-specific expression of Cre recombinase in catecholamine-containing neurons in TH-Cre KI rats.

**Figure 2.**
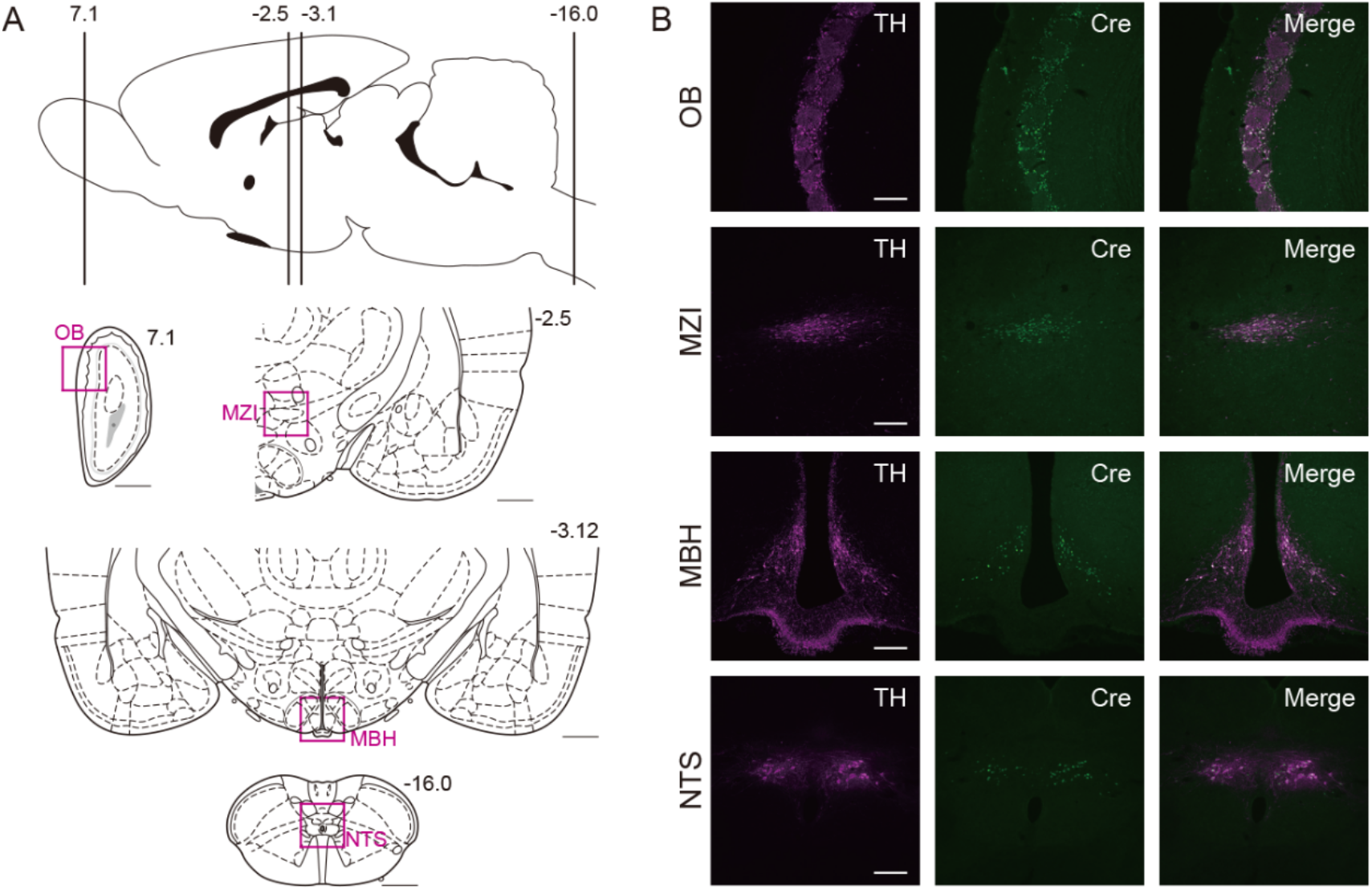
Cre transgene expression in the forebrain, hypothalamus, and posterior hindbrain of the TH-Cre KI/2-1 rat strain. (A) Representative sections from the OB, MZI, MBH, and NTS used for double fluorescent immunohistochemistry with anti-TH and anti-Cre antibodies. Anteroposterior coordinates from the bregma (mm) are shown. (B) Double immunostaining for TH and Cre. Sections from the OB, MZI, MBH, and NTS were prepared from the TH-Cre KI/2-1 strain, and doubly immunostained with two kinds of antibodies. TH-positive, Cre-positive, and merged images are shown. Scale bar: 1 mm (A), 200 µm (B).

### 3.2. Cre-loxP-mediated recombination in catecholaminergic cell types in KI rats

To confirm Cre-*lox*P-mediated recombination in catecholamine-containing neurons in TH-Cre KI/2-1 transgenic rats, we injected AAV-FLEX-GFP vector (3.7 × 10^12^ genome copies/mL) into the VTA (0.5 µL/site × 6 sites), SNc (0.5 µl/site × 4 sites) or LC (0.5 µL/site × 4 sites) of TH-Cre KI/2-1 strain. Coronal sections from the VTA, SNc, and LC were prepared from the brains, and then stained by double immunohistochemistry with anti-TH and anti-GFP antibodies (see Fig. 3 for typical images). The frequencies of Cre-*lox*P recombination in catecholamine-containing neurons, defined as the percentage of the number of TH^+^/GFP^+^ neurons divided by the total number of TH^+^ neurons, were 79 ± 1.3% for the VTA, 79 ± 3.4% for the SNc, and 90 ± 2.2% for the LC (n = 6 animals). The data indicate an efficient site-specific recombination in catecholamine-containing neurons in TH-Cre KI rats.

**Figure 3.**
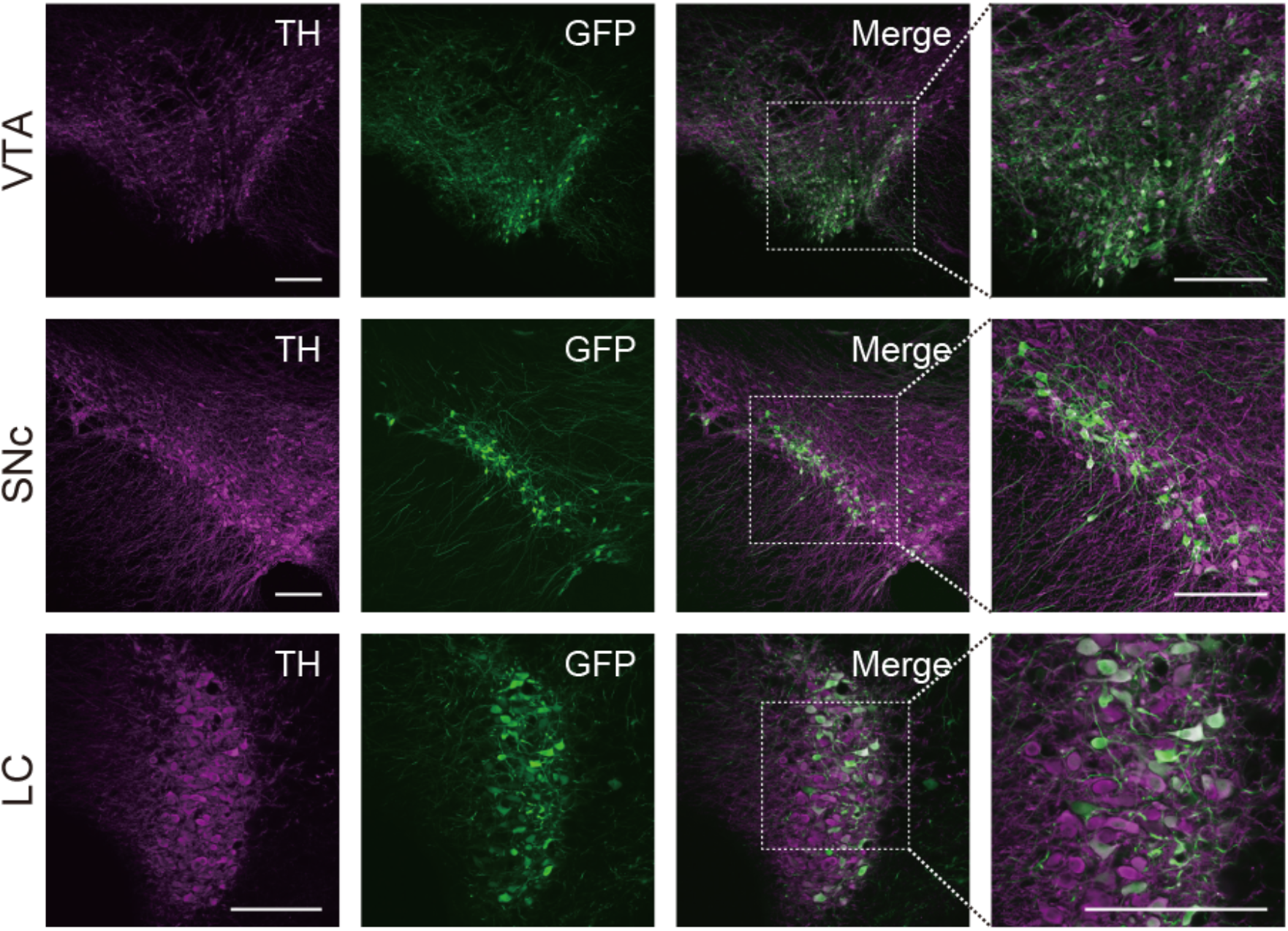
Cre-*lox*P-mediated expression of transgene in catecholaminergic cell types of the TH-Cre KI rat strain. AAV-FLEX-GFP vector was injected into the SNc, VTA, and LC of TH-Cre KI/2-1 rats, and 4 weeks later, sections from these brain regions were prepared and used for double fluorescent immunohistochemistry with anti-TH and anti-GFP antibodies. TH-positive, GFP-positive, and merged images are shown. The right photos are magnified views of the squares in the merged images. Scale bar: 200 µm.

### 3.3. TH expression level in catecholaminergic cell types in KI rats

To examine the influence of the KI mutagenesis on TH protein levels in the catecholamine-containing neurons, we performed double immunostaining for TH and MAP2 with sections from the VTA, SNc, and LC prepared from TH-Cre KI/2-1 rats and their WT littermates (see Fig 4A for representative images). For a comparison of the TH protein levels between the two kinds of rats, the immunofluorescent intensity of TH^+^ signals were normalized by the MAP2 intensity in each TH^+^ neuron. We did not find any significant differences in the TH/MAP2 ratio in the three brain regions between the genotypes (Fig 4B; unpaired *t*-tests, *t*_6_ = 1.325, *p* = 0.233 for the SNc; *t*_4.855_ = 0.550, *p* = 0.607, Levene’s test *F* = 6.418, *p* = 0.044 for the VTA; *t*_6_ = 0.589, *p* = 0.578 for the LC). Thus, the mutation in the TH-Cre KI transgenic rats did not alter TH protein levels in catecholamine-containing neurons.

**Figure 4.**
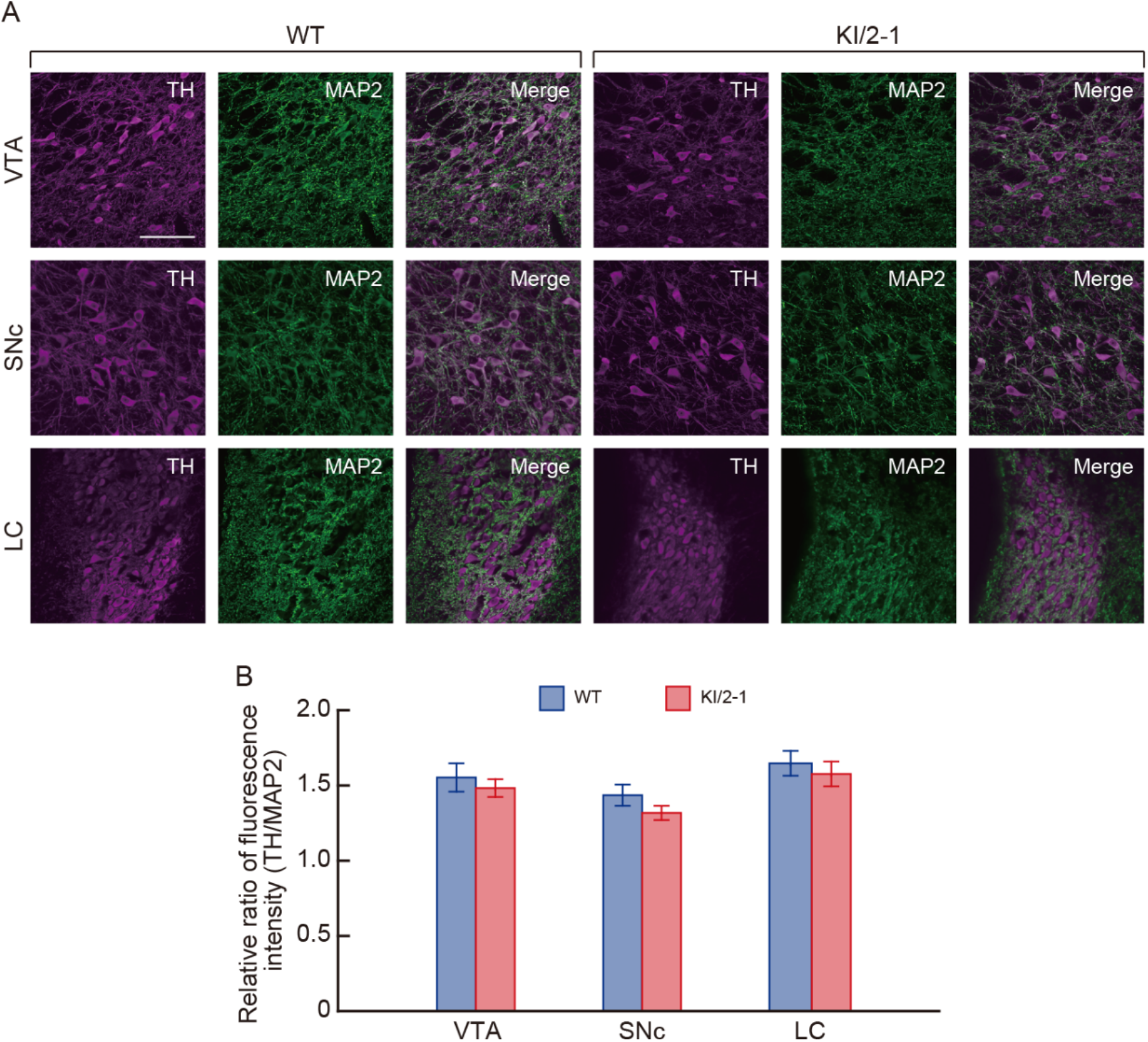
Normal TH protein level in brain regions of the TH-Cre KI/2-1 rat strain. (A) Representative sections from the VTA, SNc, and LC used for double fluorescent immunohistochemistry with anti-TH and anti-MAP2 antibodies. Sections from the VTA, SNc, and LC were prepared from the TH-Cre KI/2-1 rats and their WT littermates, and doubly immunostained with two kinds of antibodies. TH-positive, MAP2-positive, and merged images are shown. Scale bar: 100 µm. (B) Quantitation of immunofluorescent intensity of TH in each TH-positive soma located in the VTA, SNc, and LC of the TH-Cre KI/2-1 rats (*n* = 4) and their WT littermates (*n* = 4). The TH immunofluorescent intensity was normalized by using the MAP2 immunofluorescent intensity (TH/MAP2 ratio). Data are presented as mean ± *SEM*.

### 3.4. Catecholamine accumulation and behaviors in KI rats

To check the effect of the KI mutation on catecholamine accumulation, we measured DA and NE contents in the homogenates obtained from the brain regions which receive dense innervations from catecholaminergic neurons, including the ACC, NAc, and DS between the KI/2-1 rats and their WT littermates. There were no significant differences between the genotypes regarding DA contents (Fig. 5A; unpaired *t*-tests, *t*_8_ = 0.063, *p* = 0.951 for the ACC; *t*_8_ = 1.381, *p* = 0.205 for the NAc, *t*_5.659_ = 1.006, *p* = 0.356 Levene’s test *F* = 11.659, *p* = 0.009 for the DS) and NE contents (unpaired *t*-tests, *t*_8_ = 0.690, *p* = 0.510 for the ACC; *t*_8_ = 0.065, *p* = 0.949 for the NAc, *t*_8_ = 0.549, *p* = 0.598 for the DS). These results indicate that the mutation in the TH-Cre KI rats did not influence catecholamine accumulations in the brain regions.

**Figure 5.**
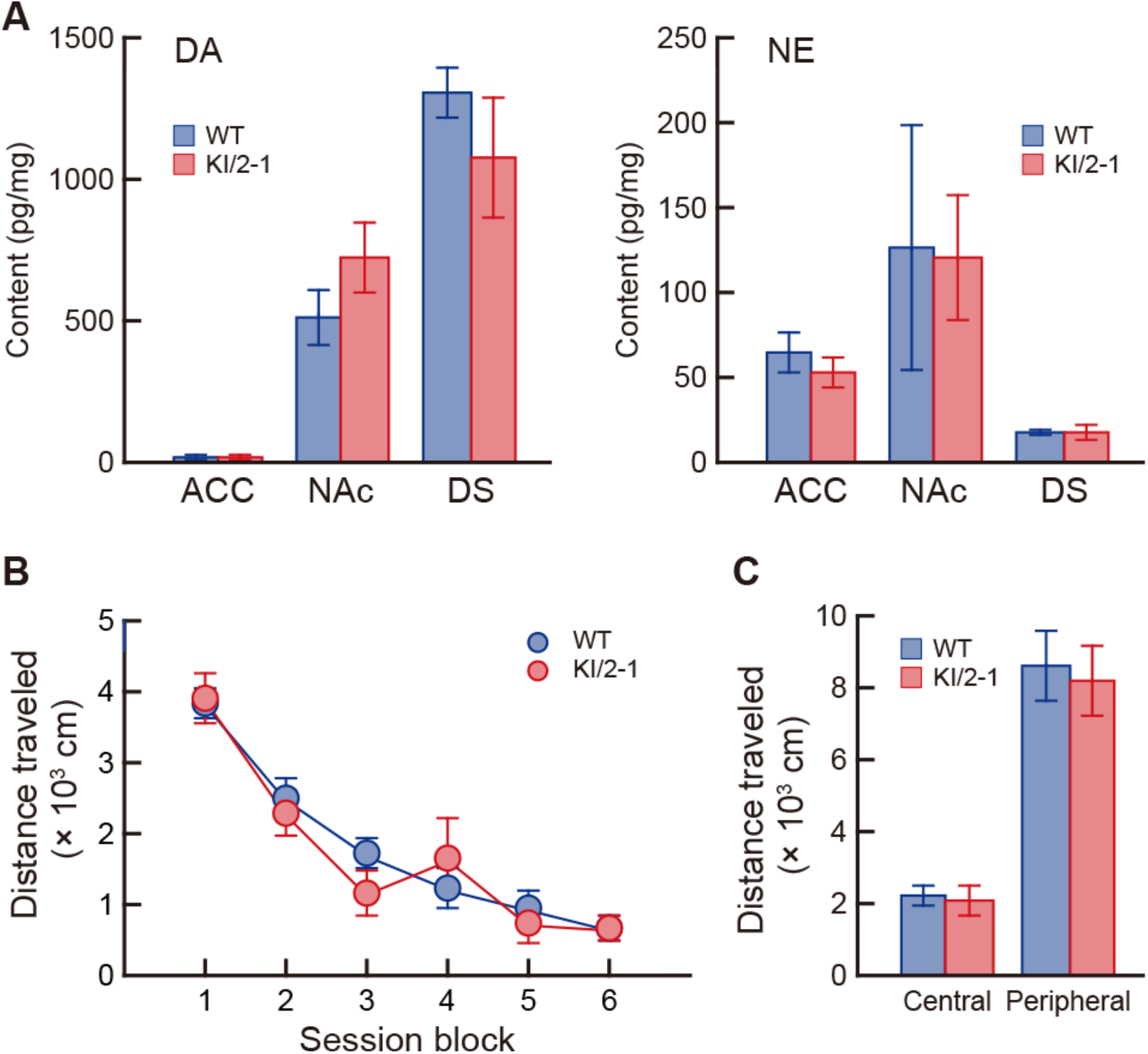
Normal catecholamine contents in the brain regions and unaffected spontaneous behaviors of TH-Cre KI/2-1 rats. (A) DA and NE accumulations in the ACC, NAc and DS of the TH-Cre KI/2-1 rats (*n* = 5) and their WT littermates (*n* = 5). The tissues of the ACC, NAc, and DS were dissected and homogenized, and catecholamine contents were determined with the HPLC system. Data are presented as mean ± *SEM*. (B) Spontaneous behaviors in a novel open field of the TH-Cre KI/2-1 rats (*n* = 5) and their WT littermates (*n* = 6). Transition of the distance traveled (in cm) of animals in a 60-min test period were plotted in blocks of 10-min. Data are presented as mean ± *SEM*. (C) Distance traveled (in cm) in the central and peripheral zones of the open field apparatus in the 60-min test period (*n* = 5 for KI/2-1, *n* = 6 for WT). Data are presented as mean ± *SEM*.

Finally, we tested the consequence of the KI mutation on the spontaneous behaviors of the rats in a novel open field. Locomotor activity was gradually decreased over the 60-min test period, and there was no significant difference between the KI/2-1 rats and their WT littermates (Fig 5B; two-way ANOVA with repeated measures: group effect *F*_1,9_ = 0.143, *p* = 0.712; time block effect *F*_5,45_ = 45.586, *p* < 0.00001; interaction *F*_5,45_ = 0.776, *p* = 0.572). In addition, when the arena was divided into the central and peripheral zones, and the distances traveled in each zone were analyzed, we did not find any significant difference between the genotypes in either zone (Fig 5C; unpaired *t*-tests, *t*_9_ = 0.239, *p* = 0.817 for the central zone; *t*_9_ = 0.323, *p* = 0.754 for the peripheral zone). The change in the time spent on the central zone of the open field is regarded as an index of anxious behavior in rodents (Coelho et al., 2014). Therefore, the TH-Cre KI transgenic rats in the present study exhibited no alterations regarding spontaneous behavior and anxiety state in the open field test.

## 4. Discussion

In our recent study, we introduced a gene cassette encoding 2A peptide connected to Cre recombinase with the nuclear localization signal into the site between the C-terminus and termination codons of rat TH open reading frame by using *Combi*-CRISPR technology, and generated two kinds of TH-Cre KI rat strains carrying an artificial insertion of a single nucleotide, A or T, into intron 12 of the TH gene (Yoshimi et al., 2021). In the present study, we first examined the expression patterns of Cre recombinase in the representative catecholaminergic cell groups in the two TH-Cre KI strains, and found similar patterns of the transgene expression, indicating no influence of a single nucleotide insertion into intron 12 on transgene expression patterns. Using one of the strains, we further found the transgene expression in other catecholamine-containing cell groups, indicating the catecholamine cell type-specific expression of Cre recombinase in our KI transgenic rats.

Based on the images obtained from double immunohistochemistry, the intracellular localization of Cre recombinase was confirmed in the nuclei in the KI rat strains. In addition, we detected many GFP-expressing cells in the representative DA and NE cell groups of the KI rats after the intracranial injection of AAV-FLEX-GFP vector, ascertaining the functional recombinase activity of Cre transgene products. In our strategy for KI mutagenesis, a precursor containing TH-2A-Cre fusion protein was synthesized and then cleaved between proline and glycine at the respective 19^th^ and 20^th^ residues in 2A peptide consisting of 21 amino acids (Kim et al., 2011). After the cleavage, two amino acids containing glycine and proline are added at the N-terminus of Cre protein with the nuclear localization signal (see Supplementary Fig. 1B). Our observations suggest that this modification does not appear to alter the intracellular localization and enzymatic activity of Cre recombinase.

Secondly, we examined TH expression and catecholamine accumulation levels in some brain regions of the TH-Cre KI rats. Based on double immunostaining, TH protein levels were apparently normal in the VTA, SNc, and LC in TH-Cre KI rats as compared to the WT animals. TH immunoreactivity was detected in the soma, dendrites, and axons in the KI strain rats, showing the normal intracellular distribution of the protein. Catecholamine analysis also indicated unaffected contents of DA and NE in the ACC, NAc, and DS, which receive abundant innervations of catecholaminergic fibers, in the KI rat strains, being consistent with the results that TH protein levels were normal in the brain regions in the strains. The cleavage of 2A peptide causes an addition of 19 extra amino acids in the C-terminus of TH protein (see Supplementary Fig. 1B). Our results suggest that the addition of extra amino acids into the C-terminus does not seem to affect the expression level, intracellular localization, or enzymatic activity of TH protein.

The activity of midbrain DA neurons is known to be involved in spontaneous locomotor activity (Boekhoudt et al., 2016; Koob et al., 1981; Pijnenburg et al., 1975). LC NE activity is reported to modulate anxiety-like behavior (Koob, 1999; McCall et al., 2015; McCall et al., 2017). In the present study, the distance travelled in an open field was gradually reduced in TH-Cre KI rats, which was similar to the WT animals, and the distances travelled in the central and peripheral zones were not significantly different between the groups, indicating the normal spontaneous and anxiety-like behaviors in the TH-Cre KI rat strains. These results suggest that the activity of catecholamine-containing neurons was not changed by the KI mutation. The results were also supported by normal catecholamine accumulation in the brain regions in the KI rat strain.

These findings suggest that the introduction of 2A peptide connected to Cre recombinase with the nuclear localization into the C-terminus of the TH open reading frame using the *Combi*-CRISPR technology achieved a useful strategy to express the functionally active recombinase in a cell type-specific manner with no significant effect on endogenous protein levels, which was accompanied by no changes in the catecholamine accumulation in the brain regions or the general behavior of the mutants. A previous study reported other KI rats designed to express Cre in a bicistronic manner with the IRES under the control of the TH gene promoter, which were generated by genomic editing using the zinc finger nuclease (Brown et al., 2013). Although these KI rats showed sufficient Cre expression to cleave the floxed DNA sequence, the targeted mutation resulted in a moderate decrease in TH mRNA and protein levels in the adrenal gland and brain, suggesting that the insertion of the IRES interferes partially the expression of the endogenous gene product (Brown et al., 2013). Comparison of biochemical impacts between the two KI strategies with the 2A peptide-mediated protein precursor and IRES-mediated bicistronic systems to express Cre recombinase suggests that the use of the 2A peptide precursor appears to have a smaller effect on the expression level of endogenous gene products relative to the introduction of the IRES into the targeted locus.

In conclusion, the mutagenesis in TH-Cre KI transgenic rats established herein does not influence catecholamine metabolism or the general behavior of animals, with no change in endogenous TH activity. This system will provide an advantageous experimental tool to investigate the molecular and cellular mechanisms underlying the morphogenesis, physiology, and pathology of catecholamine-containing neurons in the brain.

## CRediT authorship contribution statement

**Natsuki Matsushita:** Conceptualization, Methodology, Investigation, Formal analysis, Visualization, Resources, Writing - original draft, Writing - review & editing. **Kayo Nishizawa:** Conceptualization, Formal analysis, Visualization. **Shigeki Kato:** Methodology, Investigation, Formal analysis, Visualization, Writing - original draft, Writing - review & editing, Funding acquisition. **Yoshio Iguchi:** Methodology, Investigation, Formal analysis, Visualization, Writing - original draft, Writing - review & editing. **Ryoji Fukabori:** Methodology, Investigation, Formal analysis, Visualization. **Kosei Takeuchi:** Methodology, Investigation, Supervision. **Yoshiki Miyasaka:** Resources. **Tomoji Mashimo:** Resources. **Kazuto Kobayashi:** Conceptualization, Methodology, Investigation, Writing - original draft, Writing - review & editing, Supervision, Funding acquisition.

## Acknowledgements

This work was supported by grants-in-aid for Scientific Research on Transformative Research Areas (A) Adaptive Circuit Census (21H05244) to K.K from the Ministry of Education, Science, Sports, and Culture of Japan, and by a grant-in-aid for Young Scientists (A) (25702053) from the Japan Society for the Promotion of Science (S.K.). We are grateful to H. Hashimoto, M. Kikuchi, and Y. Nakazato for technical support during animal experiments, and to T. Kobayashi for helpful illustrations.

## Data availability

The data that support the findings of this study are publicly available in Mendeley Data (https://doi.org/10.17632/3k99jn38w3.1).

## Abbreviations

DA: dopamine
EPI: epinephrine
KI: knock-in
LC: locus coeruleus
NE: norepinephrine
PBS: phosphate-buffered saline
SNc: substantia nigra pars compacta
TH: tyrosine hydroxylase
VTA: ventral tegmental area

## Figures and Figure legends

**Supplementary Fig. 1.**
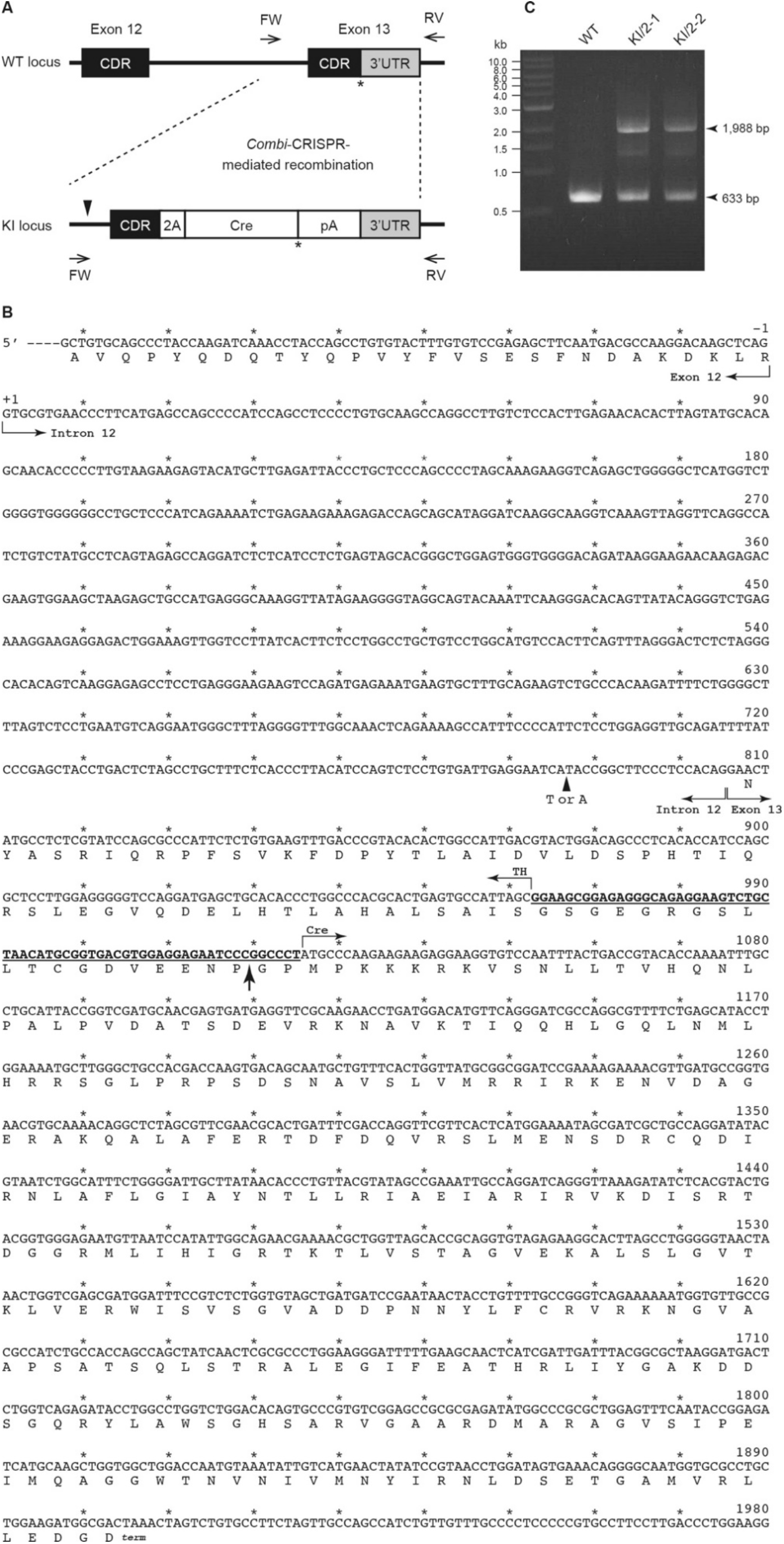
TH-Cre KI transgenic rats. (A) Structure of WT and KI alleles of rat TH locus. A gene cassette encoding 2A peptide fused to Cre recombinase with nuclear localization signal followed by the polyadenylation signal (pA) from the bovine growth hormone gene is introduced between the C-terminus of protein-coding region (CDR) and 3’-untranslated region (3’-UTR) in exon 13 of the rat TH gene by the *Combi*-CRISPR technology. The asterisks show the position of stop codon, and the arrowhead indicates the location of a single nucleotide insertion (T or A) in the KI allele. The arrows represent the position and direction of forward (FW) and reverse (RV) primers for PCR amplification. (B) Partial nucleotide sequence from exon 12 to 3’-UTR of the TH-Cre KI alleles. The nucleotides are numbered from the first nucleotide in intron 12 as +1. Amino acids are shown beneath the corresponding nucleotide sequences. The arrowheads indicate the position of a single nucleotide insertion in the KI allele: T for KI/2-1 strain and A for KI/2-2 strain. Nucleotide sequences corresponding to 2A peptide are underlined and the cleavage site is shown by the arrow. (C) Genotyping of the offspring by PCR amplification with genomic DNA prepared from tail clips. Agarose gel electrophoresis indicates DNA fragments of 633 bp and 1,988 bp derived from WT and KI alleles, respectively. Sizes of molecular markers (kb) are shown on the left of the image.

## References

Arenas, E., Denham, M., Villaescusa, JC., 2015. How to make a midbrain dopaminergic neuron. Development 142 (11): 1918–36. doi: 10.1242/dev.097394

Berardelli, A., Wenning, GK., Antonini, A., Berg, D., Bloem, BR., Bonifati, V., Brooks, D., Burn, D.J., Colosimo, C., Fanciulli, A., Ferreira, J., Gasser, T., Grandas, F., Kanovsky, P., Kostic, V., Kulisevsky, J., Oertel, W., Poewe, W., Reese, JP., Relja, M., Ruzicka, E., Schrag, A., Seppi, K., Taba, P., Vidailhet, M., 2013. EFNS/MDS-ES/ENS [corrected] recommendations for the diagnosis of Parkinson’s disease. Eur. J. Neurol. 20 (1): 16–34. doi: 10.1111/ene.12022

Björklund, A., Dunnett, SB., 2007. Dopamine neuron systems in the brain: an update. Trends. Neurosci. 30 (5): 194–202. doi: 10.1016/j.tins.2007.03.006

Boekhoudt, L., Omrani, A., Luijendijk, MC., Wolterink-Donselaar, IG., Wijbrans, EC., van der Plasse, G., Adan, RA., 2016. Chemogenetic activation of dopamine neurons in the ventral tegmental area, but not substantia nigra, induces hyperactivity in rats. Eur. Neuropsychopharmacol. 26 (11): 1784–93. doi: 10.1016/j.euroneuro.2016.09.003

Brown, AJ., Fisher, DA., Kouranova, E., McCoy, A., Forbes, K., Wu, Y., Henry R., Ji, D., Chambers, A., Warren, J., Shu, W., Weinstein, EJ., Cui, X., 2013. Whole-rat conditional gene knockout via genome editing. Nat. Methods 10 (7): 638–40. doi: 10.1038/nmeth.2516

Brown, ER., Coker, GT., 3rd, O’Malley, KL., 1987. Organization and evolution of the rat tyrosine hydroxylase gene. Biochemistry 26 (16): 5208–12. doi: 10.1021/bi00390a046

Buttery, PC., Barker, RA., 2020. Gene and cell-based therapies for Parkinson’s disease: Where are we? Neurotherapeutics 17 (4): 1539–62. doi: 10.1007/s13311-020-00940-4

Cassidy, CM., Balsam, PD., Weinstein, JJ., Rosengard, RJ., Slifstein, M., Daw, ND., Abi-Dargham, A., Horga, G., 2018. A perceptual inference mechanism for hallucinations linked to striatal dopamine. Curr. Biol. 28 (4): 503–14.e4. doi: 10.1016/j.cub.2017.12.059

Coelho, JE., Alves, P., Canas, PM., Valadas, JS., Shmidt, T., Batalha, VL., Ferreira, DG., Ribeiro, JA., Bader, M., Cunha, RA., do Couto, FS., Lopes, LV., 2014. Overexpression of adenosine A2A receptors in rats: Effects on depression, locomotion, and anxiety. Front. Psychiatry 5:67. doi: 10.3389/fpsyt.2014.00067

Gelman, DM., Noaín, D., Avale, ME., Otero, V., Low, MJ., Rubinstein, M., 2003. Transgenic mice engineered to target Cre/loxP-mediated DNA recombination into catecholaminergic neurons. Genesis 36 (4): 196–202. doi: 10.1002/gene.10217

Gerfen, CR., Paletzki, R., Heintz, N., 2013. GENSAT BAC cre-recombinase driver lines to study the functional organization of cerebral cortical and basal ganglia circuits. Neuron 80 (6): 1368–83. doi: 10.1016/j.neuron.2013.10.016

Gong, S., Doughty, M., Harbaugh, CR., Cummins, A., Hatten, ME., Heintz, N., Gerfen, CR., 2007. Targeting Cre recombinase to specific neuron populations with bacterial artificial chromosome constructs. J. Neurosci. 27 (37): 9817–23. doi: 10.1523/JNEUROSCI.2707-07.2007

Gordon, DF., Quick, DP., Erwin, CR., Donelson, JE., Maurer, RA., 1983. Nucleotide sequence of the bovine growth hormone chromosomal gene. Mol. Cell. Endocrinol. 33 (1): 81–95. doi: 10.1016/0303-7207(83)90058-8

Hennecke, M., Kwissa, M., Metzger, K., Oumard, A., Kröger, A., Schirmbeck, R., Reimann, J., Hauser, H., 2001. Composition and arrangement of genes define the strength of IRES-driven translation in bicistronic mRNAs. Nucleic Acids Res. 29 (16): 3327–34. doi: 10.1093/nar/29.16.3327.

Hirsch, MR., Tiveron, MC., Guillemot, F., Brunet, JF., Goridis, C., 1998. Control of noradrenergic differentiation and Phox2a expression by MASH1 in the central and peripheral nervous system. Development 125 (4): 599–608. doi: 10.1242/dev.125.4.599

Holm, PC., Akerud, P., Wagner, J., Arenas, E., 2002. Neurturin is a neuritogenic but not a survival factor for developing and adult central noradrenergic neurons. J. Neurochem. 81 (6): 1318–27. doi: 10.1046/j.1471-4159.2002.00926.x

Inouye, S., Tsuji, FI., 1994. Aequorea green fluorescent protein. Expression of the gene and fluorescence characteristics of the recombinant protein. FEBS Lett. 341 (2-3): 277–80. doi: 10.1016/0014-5793(94)80472-9

Inui, M., Miyado, M., Igarashi, M., Tamano, M., Kubo, A., Yamashita, S., Asahara, H., Fukami, M., Takada, S., 2014. Rapid generation of mouse models with defined point mutations by the CRISPR/Cas9 system. Sci. Rep. 4:5396. doi: 10.1038/srep05396

Jaumotte, JD., Saarma, M., Zigmond, MJ., 2021. Protection of dopamine neurons by CDNF and neurturin variant N4 against MPP+ in dissociated cultures from rat mesencephalon. PLoS One 16:e0245663. doi: 10.1371/journal.pone.0245663

Kato, S., Fukabori, R., Nishizawa, K., Okada, K., Yoshioka, N., Sugawara, M., Maejima, Y., Shimomura, K., Okamoto, M., Eifuku, S., Kobayashi, K., 2018. Action selection and flexible switching controlled by the intralaminar thalamic neurons. Cell Rep. 22:2370–82. doi: 10.1016/j.celrep.2018.02.016

Kim, JH., Lee, SR., Li, LH., Park, HJ., Park, JH., Lee, KY., Kim, MK., Shin, BA., Choi, SY., 2011. High cleavage efficiency of a 2A peptide derived from porcine teschovirus-1 in human cell lines, zebrafish and mice. PLoS One 6:e18556. doi: 10.1371/journal.pone.0018556

Kobayashi, K., Morita, S., Mizuguchi, T., Sawada, H., Yamada, K., Nagatsu, I., Fujita, K., Nagatsu, T., 1994. Functional and high level expression of human dopamine beta-hydroxylase in transgenic mice. J. Biol. Chem. 269 (47): 29725–31. doi: 10.1016/S0021-9258(18)43941-5

Koob, GF., 1999. Corticotropin-releasing factor, norepinephrine, and stress. Biol. Psychiatry 46 (9): 1167–80. doi: 10.1016/s0006-3223(99)00164-x

Koob, GF., Stinus, L., Le Moal, M., 1981. Hyperactivity and hypoactivity produced by lesions to the mesolimbic dopamine system. Behav. Brain Res. 3 (3): 341–59. doi: 10.1016/0166-4328(81)90004-8

Kumar, A., Kopra, J., Varendi, K., Porokuokka, LL., Panhelainen, A., Kuure, S., Marshall, P., Karalija, N., Härma, MA., Vilenius, C., Lilleväli, K., Tekko, T., Mijatovic, J., Pulkkinen, N., Jakobson, M., Jakobson, M., Ola, R., Palm, E., Lindahl, M., Strömberg, I., Võikar, V., Piepponen, TP., Saarma, M., Andressoo, JO., 2015. GDNF Overexpression from the native locus reveals its role in the nigrostriatal dopaminergic system function. PLoS Genet. 11 (12): e1005710. doi: 10.1371/journal.pgen.1005710

Lindeberg, J., Usoskin, D., Bengtsson, H., Gustafsson, A., Kylberg, A., Söderström, S., Ebendal, T., 2004. Transgenic expression of Cre recombinase from the tyrosine hydroxylase locus. Genesis 40 (2): 67–73. doi: 10.1002/gene.20065

Łobocka, MB., Rose, DJ., Plunkett, G., 3rd, Rusin, M., Samojedny, A., Lehnherr, H., Yarmolinsky, MB., Blattner, FR., 2004. Genome of bacteriophage P1. J. Bacteriol. 186 (21): 7032–68. doi: 10.1128/jb.186.21.7032-7068.2004

Mäki-Marttunen, V., Andreassen, OA., Espeseth, T., 2020. The role of norepinephrine in the pathophysiology of schizophrenia. Neurosci. Biobehav. Rev.118: 298–314. doi: 10.1016/j.neubiorev.2020.07.038

Matsushita, N., Kobayashi, K., Miyazaki, J., Kobayashi, K., 2004. Fate of transient catecholaminergic cell types revealed by site-specific recombination in transgenic mice. J. Neurosci. Res. 78 (1): 7–15. doi: 10.1002/jnr.20229

Matsushita, N., Kato, S., Nishizawa, K., Sugawara, M., Takeuchi, K., Miyasaka, Y., Mashimo, T., Kobayashi, K., 2022. Highly selective transgene expression through double-floxed inverted orientation system by using a unilateral spacer sequence. bioRxiv. doi: 10.1011/2022.04.11.487972

McCall, JG., Al-Hasani, R., Siuda, ER., Hong, DY., Norris, AJ., Ford, CP., Bruchas, MR., 2015. CRH engagement of the locus coeruleus noradrenergic system mediates stress-induced anxiety. Neuron 87 (3): 605–20. doi: 10.1016/j.neuron.2015.07.002

McCall, JG., Siuda, ER., Bhatti, DL., Lawson, LA., McElligott, ZA., Stuber, GD., Bruchas, MR., 2017. Locus coeruleus to basolateral amygdala noradrenergic projections promote anxiety-like behavior. eLife 6:e18247. doi: 10.7554/eLife.18247

Miura, H., Gurumurthy, CB., Sato, T., Sato, M., Ohtsuka, M., 2015. CRISPR/Cas9-based generation of knockdown mice by intronic insertion of artificial microRNA using longer single-stranded DNA. Sci. Rep. 5:12799. doi: 10.1038/srep12799

Morin, X., Cremer, H., Hirsch, MR., Kapur, RP., Goridis, C., Brunet, JF., 1997. Defects in sensory and autonomic ganglia and absence of locus coeruleus in mice deficient for the homeobox gene Phox2a. Neuron 18 (3): 411–23. doi: 10.1016/s0896-6273(00)81242-8

Nestler, EJ., Lüscher, C., 2019. The molecular basis of drug addiction: linking epigenetic to synaptic and circuit mechanisms. Neuron 102 (1): 48–59. doi: 10.1016/j.neuron.2019.01.016

Palasz, E., Wysocka, A., Gasiorowska, A., Chalimoniuk, M., Niewiadomski, W., Niewiadomska, G., 2020. BDNF as a promising therapeutic agent in Parkinson’s disease. Int. J. Mol. Sci. 21 (3): 1170. doi: 10.3390/ijms21031170

Pattyn, A., Goridis, C., Brunet, JF., 2000. Specification of the central noradrenergic phenotype by the homeobox gene Phox2b. Mol. Cell. Neurosci. 15 (3): 235–43. doi: 10.1006/mcne.1999.0826

Paxinos, G., Watson, C., 2007. The Rat Brain in Stereotaxic Coordinates, Ed 6. Sydney: Academic Press.

Pijnenburg, AJ., Honig, WM., Van Rossum, JM., 1975. Inhibition of d-amphetamine-induced locomotor activity by injection of haloperidol into the nucleus accumbens of the rat. Psychopharmacologia 41 (2): 87–95. doi: 10.1007/bf00421062

Quadros, RM., Miura, H., Harms, DW., Akatsuka, H., Sato, T., Aida, T., Redder, R., Richardson, GP., Inagaki, Y., Sakai, D., Buckley, SM., Seshacharyulu, P., Batra, SK., Behlke, MA., Zeiner, SA., Jacobi, AM., Izu, Y., Thoreson, WB., Urness, LD., Mansour, SL., Ohtsuka, M., Gurumurthy, CB., 2017. Easi-CRISPR: a robust method for one-step generation of mice carrying conditional and insertion alleles using long ssDNA donors and CRISPR ribonucleoproteins. Genome Biol. 18 (1): 92. doi: 10.1186/s13059-017-1220-4

Ranjbar-Slamloo, Y., Fazlali, Z., 2020. Dopamine and noradrenaline in the brain; overlapping or dissociate functions? Front. Mol. Neurosci. 12:334. doi: 10.3389/fnmol.2019.00334

Reiriz, J., Holm, PC., Alberch, J., Arenas, E., 2002. BMP-2 and cAMP elevation confer locus coeruleus neurons responsiveness to multiple neurotrophic factors. J. Neurobiol. 50 (4): 291–304. doi: 10.1002/neu.10034

Robbins, TW., Everitt, BJ., 1982. Functional studies of the central catecholamines. Int. Rev. Neurobiol. 23:303–65. doi: 10.1016/s0074-7742(08)60628-5

Robertson, SD., Plummer, NW., de Marchena, J., Jensen, P., 2013. Developmental origins of central norepinephrine neuron diversity. Nat. Neurosci. 16 (8): 1016–23. doi: 10.1038/nn.3458

Ross, JA., Van Bockstaele, EJ., 2021. The locus coeruleus-norepinephrine system in stress and arousal: unraveling historical, current, and future perspectives. Front. Psychiatry 11:601519. doi: 10.3389/fpsyt.2020.601519

Schapira, AH., Emre, M., Jenner, P., Poewe, W., 2009. Levodopa in the treatment of Parkinson’s disease. Eur. J. Neurol. 16 (9): 982–9. doi: 10.1111/j.1468-1331.2009.02697.x

Schindelin, J., Arganda-Carreras, I., Frise, E., Kaynig, V., Longair, M., Pietzsch, T., Preibisch, S., Rueden, C., Saalfeld, S., Schmid, B., Tinevez, JY., White, DJ., Hartenstein, V., Eliceiri, K., Tomancak, P., Cardona, A., 2012. Fiji: an open-source platform for biological-image analysis. Nat. Methods 9 (7): 676–82. doi: 10.1038/nmeth.2019

Silvetti, M., Wiersema, JR., Sonuga-Barke, E., Verguts, T., 2013. Deficient reinforcement learning in medial frontal cortex as a model of dopamine-related motivational deficits in ADHD. Neural Netw. 46:199–209. doi: 10.1016/j.neunet.2013.05.008

Takai, A., Nakano, M., Saito, K., Haruno, R., Watanabe, TM., Ohyanagi, T., Jin, T., Okada, Y., Nagai, T., 2015. Expanded palette of Nano-lanterns for real-time multicolor luminescence imaging. Proc. Natl. Acad. Sci. U. S. A. 112 (14): 4352–6. doi: 10.1073/pnas.1418468112

Vogel-Höpker, A., Rohrer, H., 2002. The specification of noradrenergic locus coeruleus (LC) neurones depends on bone morphogenetic proteins (BMPs). Development 129 (4): 983–91. doi: 10.1242/dev.129.4.983

Wang, M., Ling, KH., Tan, JJ., Lu, CB., 2020. Development and differentiation of midbrain dopaminergic neuron: from bench to bedside. Cells 9 (6): 1489. doi: 10.3390/cells9061489

Witten, IB., Steinberg, EE., Lee, SY., Davidson, TJ., Zalocusky, KA., Brodsky, M., Yizhar, O., Cho, SL, Gong, S., Ramakrishnan, C., Stuber, GD., Tye, KM., Janak, PH., Deisseroth, K., 2011. Recombinase-driver rat lines: tools, techniques, and optogenetic application to dopamine-mediated reinforcement. Neuron 72 (5): 721–33. doi: 10.1016/j.neuron.2011.10.028

Yang, H., Wang, H., Shivalila, CS., Cheng, AW., Shi, L., Jaenisch, R., 2013. One-step generation of mice carrying reporter and conditional alleles by CRISPR/Cas-mediated genome engineering. Cell 154 (6): 1370–9. doi: 10.1016/j.cell.2013.08.022

Yoshimi, K., Oka, Y., Miyasaka, Y., Kotani, Y., Yasumura, M., Uno, Y., Hattori, K., Tanigawa, A., Sato, M., Oya, M., Nakamura, K., Matsushita, N., Kobayashi, K., Mashimo, T., 2021. Combi-CRISPR: combination of NHEJ and HDR provides efficient and precise plasmid-based knock-ins in mice and rats. Hum. Genet. 140 (2): 277–87. doi: 10.1007/s00439-020-02198-4

Zhou, Y., Aran, J., Gottesman, MM., Pastan I, 1998. Co-expression of human adenosine deaminase and multidrug resistance using a bicistronic retroviral vector. Hum. Gene Ther. 9 (3): 287–93. doi: 10.1089/hum.1998.9.3-287

